# The RacGAP βChimaerin is essential for cerebellar granule cell migration

**DOI:** 10.1101/164897

**Authors:** Jason A. Estep, Wenny Wong, Yiu-Cheung E. Wong, Brian M. Loui, Martin M. Riccomagno

## Abstract

During mammalian cerebellar development, postnatal granule cell progenitors proliferate in the outer part of the External Granule Layer (EGL). Postmitotic granule progenitors migrate tangentially in the inner EGL before switching to migrate radially inward, past the Purkinje cell layer, to achieve their final position in the mature Granule Cell Layer (GCL). Here, we show that the RacGAP β-chimaerin is expressed by a small population of late-born, premigratory granule cells. β-chimaerin deficiency causes a subset of granule cells to become arrested in the EGL, where they differentiate and form ectopic neuronal clusters. These clusters of granule cells are able to recruit aberrantly projecting mossy fibers. Collectively, these data suggest a role for β-chimaerin as an intracellular mediator of Cerebellar Granule Cell radial migration.

## Introduction

Proper morphogenesis of the vertebrate Central Nervous System (CNS) relies on the tight spatiotemporal control of cell proliferation, differentiation, migration and guidance events. In the mammalian cerebellum, Granule Cells (GCs) undergo a prolonged and highly stereotyped migration that begins embryonically and completes late postnatally ^1^. In the mouse, beginning at embryonic day 12 (E12), granule cell precursors (GCPs) are born from the rhombic lip and migrate tangentially to cover the cerebellar anlage ^2^, forming a secondary germinal zone, the External Granule Layer (EGL). Postnatally, GPCs in the EGL exit the cell cycle and travel inwards, splitting the EGL into an upper, mitotically active (outer EGL, oEGL) and a lower, migratory layer (inner EGL, iEGL) (Fig. 1a). These postmitotic GCPs grow two horizontal processes and migrate tangentially in all directions, before growing a third perpendicular leading process. Using this leading process GCPs migrate radially inward along Bergmann Glial fibers, past the Purkinje Cell (PC) Layer, to occupy their final location in the mature Granule Cell Layer (GCL) ^3,4^ Cerebellar GC migration has been shown to be influenced by a wide set of guidance cues, including the chemokine SDF-1 ^5^, Slit2/Robos ^6^, Plexins/Semaphorins ^7-9^, brain-derived neurotrophic factor (BDNF) ^10^, Vascular Endothelial Growth Factor (VEGF) ^11^, and others. However, the cytosolic machinery responsible for effecting and directing the cellular response downstream of these ligand-receptor pairs remains largely unexplored.

**Figure 1:**
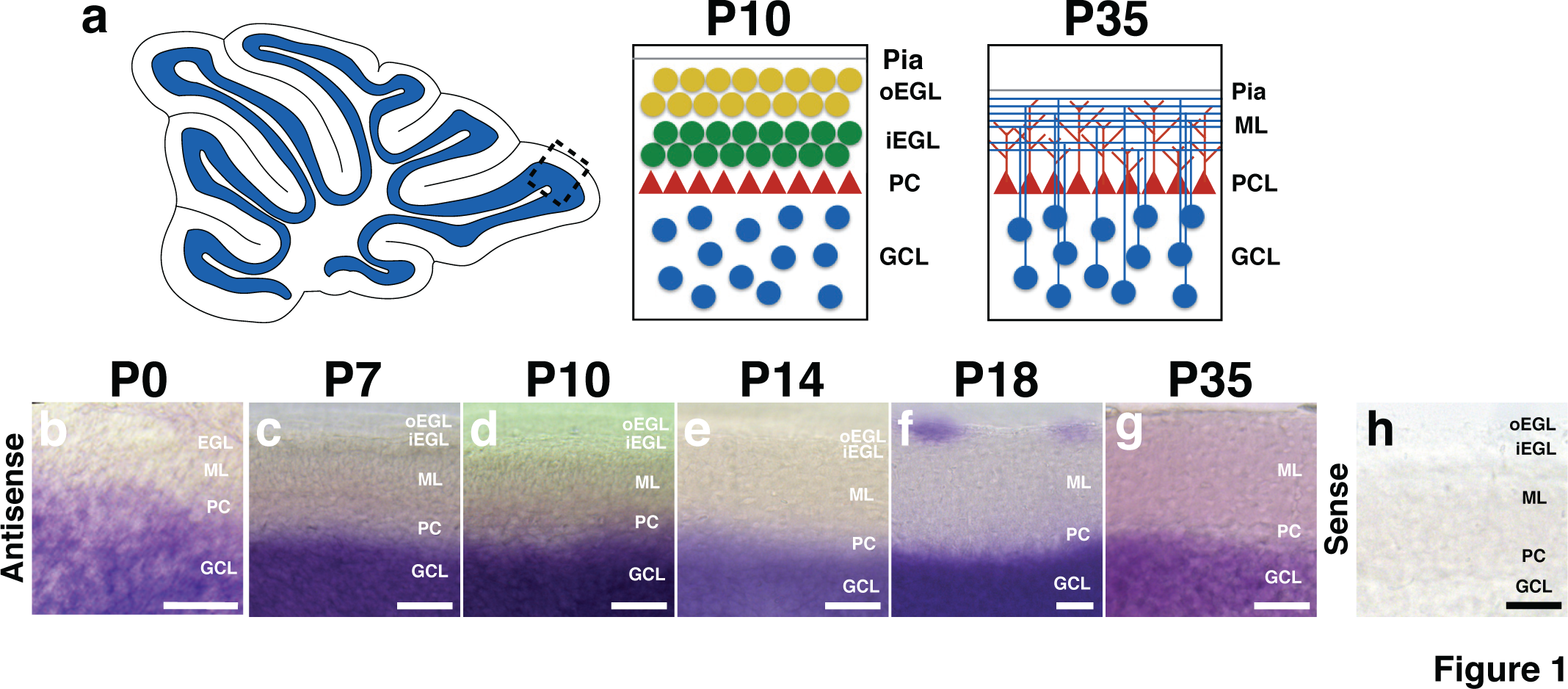
β-chimaerin expression in the postnatal cerebellum. **(a)**Developmental maturation of cerebellar granule cells. At early postnatal stages, mitotically active granule cell precursors (GCPs, yellow) populate the outer External Granule Layer (EGL). Postmitotic granule cell precursors (green) move to the inner EGL, where they grow two horizontal processes and migrate tangentially to expand across the surface of the cerebellum. These cells eventually grow a third perpendicular process and begin migrating radially inward along Bergmann glial fibers, past the Purkinje Cell layer (PCL, red triangles), to form the mature Granule Cell Layer (GCL). Mature granule cells (blue) extend their axons back to the Molecular Layer (ML) to produce parallel fibers that provide Glutamatergic inputs on Purkinje Cell dendrites. (**b-h**) *In situ hybridization* in C57/BL6J mice using a probe against β-chimaerin *(Chn2)* transcript. *Chn2* shows robust expression in the GCL at all postnatal stages. Notably, we detected expression in the EGL at P18, but this expression did not persist in adult (P35) animals. Hybridization with a sense probe does not result in any detectable signal at any of these stages (P14 is shown in h). Scale bar, 50μm for all.

The Rho family of small G-Proteins, or GTPases, plays essential roles in vertebrate CNS development, influencing a wide range of developmental processes, including cell migration, cell polarity, axon pathfinding, and dendritic remodeling through their ability to modulate cytoskeletal structure ^12,13^. GTPases exists in two states: an active GTP-bound state and inactive GDP-bound state ^14^ Precise subcellular regulation of GTPase activity is essential in maintaining proper cellular function, and neurons achieve this using positive regulators, Rho Guanine Nucleotide Exchange Factors (or RhoGEFs) and negative regulators, Rho GTPase Activating Proteins (or RhoGAPs) ^14,15^. Disruption of RhoGTPase activity or their regulators’ function has been associated with a broad array of behavioral and developmental disorders ^15,16^. The chimaerin family of RhoGAPs consists of two genes: α-chimaerin (*CHN1*)and β-chimaerin (*CHN2*). They posses specific GAP activity toward Rac family GTPases, which are key modulators of actin filaments ^17^ In neural development, α-chimaerin has been shown to play roles in Ephrin-mediated circuit formation ^18-21^, cortical migration ^22^, optic tract axon guidance ^23,24,^, and hippocampal dendritic arbor pruning ^25^. The *in vivo* role of β-chimaerin in neural development was unexplored until recently, where it was shown to effect hippocampal dentate gyrus axon pruning by regulating Rac1 activity downstream of Sema3F/Neuropilin-2 signaling ^26^. Of note, β-chimaerin has been shown to be strongly expressed in GCs in the adult ^27^, but its function during cerebellar morphogenesis is unknown. Here, we show a functional requirement for β-chimaerin during cerebellar development. We find that β-chimaerin is necessary for a small subset of granule cells to complete their migratory route from the EGL to the GCL.

## Results

### β-chimaerin is specifically expressed in the Granule Cell Layer of the mouse cerebellum

β-chimaerin has been previously shown to be expressed in the adult cerebellum ^27^ To explore the developmental expression profile of β-chimaerin in the cerebellum, we performed *in situ hybridization* in *C57/BL6J* mice to visualize β-chimaerin *(Chn2)* messenger RNA (mRNA) at several postnatal stages (Fig. 1b-h). We found *Chn2* mRNA was strongly expressed in the GCL at all the postnatal ages tested. Interestingly, we observed *Chn2* expression in small clusters of cells in the Molecular Layer (ML) of postnatal day 18 (P18) animals (Fig. 1f). This stage represents one of the last postnatal stages before the EGL dissolves. This ML expression did not persist into adulthood, disappearing by P35 (Fig. 1g).

### β-chimaerin deficient mice display ectopic neuronal clusters on the cerebellar surface

As *Chn2* transcript was found to be robustly expressed in the cerebellum at all postnatal stages examined, we asked whether β-chimaerin played a functional role during cerebellar development. We took advantage of a previously generated knock-in mouse that expresses beta-galactosidase (βgal) from the endogenous *Chn2* locus, rendering the *Chn2* gene inactive ^26^. We generated adult (P35) mice homozygous for a *Chn2* null allele (*Chn2^−/−^*) and compared their cerebellar structure to WT (*Chn2^+/+^*) littermate controls (Fig. 2a-c). We observed no gross alterations to cerebellar lobule formation or cortical lamination in *Chn2^−/−^* mutants. However, we did see large ectopic clusters of cells aggregating in the ML of mutant animals (Fig. 2b, white arrows). These clusters strongly co-labeled with the pan-neuronal marker NeuN and an antibody raised against βgal, indicating that these clusters consist of ectopic cells that normally express *Chn2*(Fig. 2 d,e). Sparse NeuN labeling was also seen in the ML of both WT and *Chn2^−/−^* genotypes (Fig. 2d,e), and likely represents the stellate and basket cells known to occupy this region. We next asked if β-chimaerin function is required in a dose-dependent manner for normal cerebellar development. We quantified the number of neuronal ectopias in *Chn2^+/+,^ Chn2^+/−^*, *Chn2^−/−^* adult animals and found a highly significant increase in the number of ectopias in *Chn2^−/−^* mutants as compared to WT and *Chn2^+/−^* animals (p<0.01 for both comparisons) (Fig. 2c).

**Figure 2:**
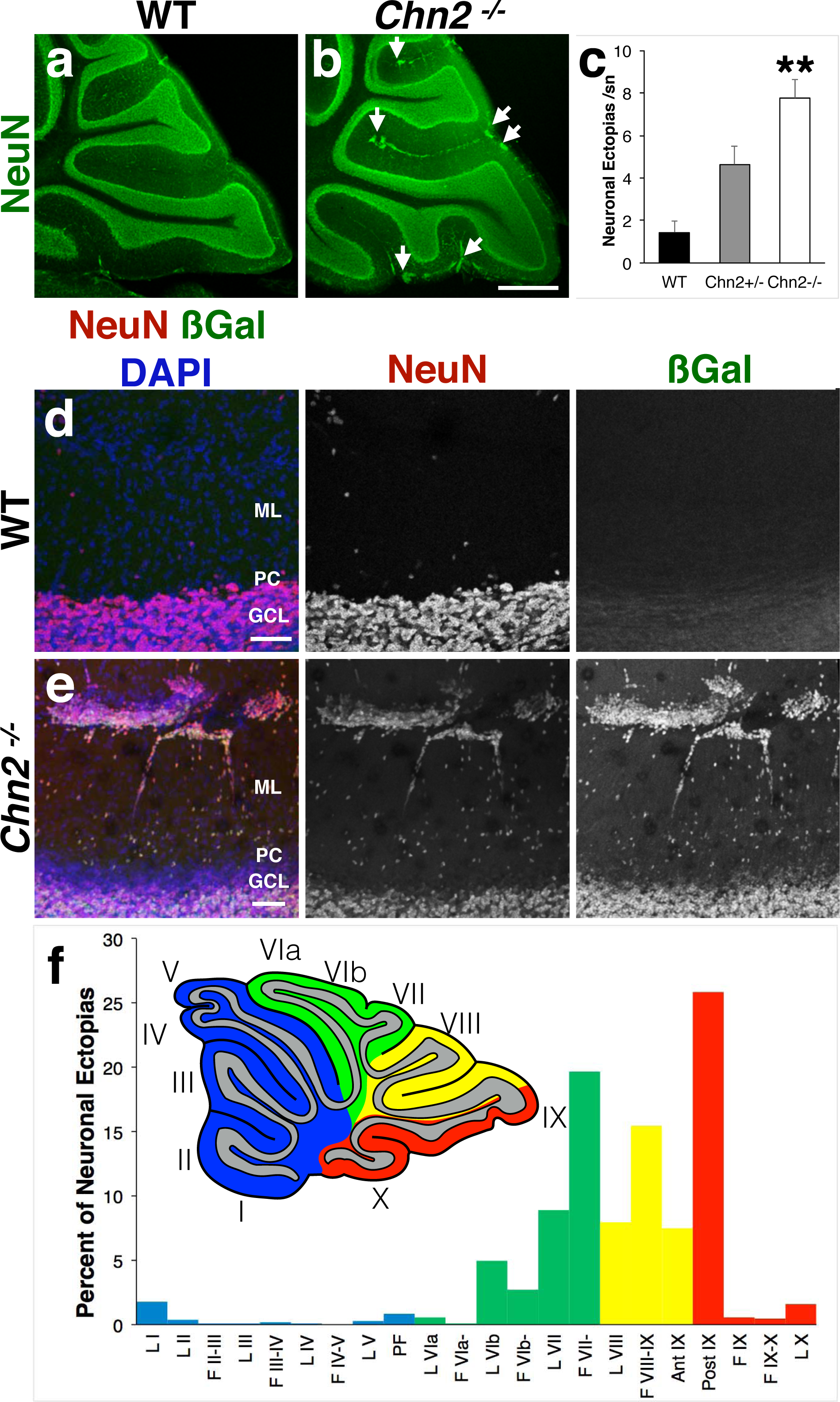
β-chimaerin deficiency causes neuronal ectopic clusters to form along the cerebellar folia in an asymmetrical pattern. **(a, b)**Immunoflourescence of the pan-neuronal marker NeuN in adult (P35) WT and *Chn2^−/−^* animals. Ectopic clusters of neurons are observed in the ML in *Chn2^−/−^* animals (white arrows), but these mutants display no other changes in overall cerebellar structure. Scale bar, 500μm. (**c**) Quantification of the average number of neuronal ectopias per 150μm section across genotypes. There is a highly significant difference among the three genotypes (n=9; One-way ANOVA, p=4.4996e-05). While there appears to be a step-wise increase in the average number of ectopias found in *Chn2^+/+^, Chn2^+/−^,* and *Chn2^−/−^* mice, only *Chn2^−/−^* show a significant increase in the frequency of ectopias as compared to *Chn2^+/+^* and *Chn2^+/-^* animals (**p<0.01 for both comparisons, Tukey HSD test). Error bars represent SEM. (**d, e**) Immunoflourescence with antibodies recognizing both NeuN and betagalactosidase (βgal) reveal that these neuronal ectopias strongly co-label with both markers. (**e**) Schematic and graph representing the percent distribution of ectopic clusters across the cerebellum in *Chn2^−/−^* animals. The cerebellum may be divided into four principle regions: Anterior (blue), Central (Green), Posterior (Yellow), and Nodal (Red); each region may be further divided into several individual folds, or Lobules (I-X). We found that ectopias most commonly occur in posterior and nodal lobules and fissures, with enrichment in the fissure separating lobules VII-VIII and on the posterior side of lobule IX (collectively accounting for 45% of all ectopias scored; n=1068 ectopias across nine animals).

The adult cerebellum can be organizationally divided into four domains: Anterior, Central, Posterior, and Nodular. Each region, in turn, is physically divided into lobules, numbered I-X in mice ^28^. Closely examining the P18 *in situ hybridization* data, we noticed that the majority of ML *Chn2* transcript expression occurred in more posterior sections, particularly Lobules VII-IX and the fissures separating them (data not shown). Therefore, we asked if the NeuN-positive clusters we observed in *Chn2^−/−^* followed a similar pattern of distribution. Indeed, we found that NeuN-positive ectopias were more prevalent in the fissure separating lobules VII and VIII and on the posterior side of lobule IX (Fig. 2f for schematic and percent distribution). These two locations collectively account for approximately 45% of all ectopic clusters scored (n=1068 ectopias across nine *Chn2^−/−^* animals). Collectively, these data suggest that *Chn2* is expressed by a small subset of late radially migrating neurons prior to their arrival to the GCL, and that loss of β-chimaerin function causes these cells to fully arrest in the EGL.

### The ectopic clusters contain mature granule cells, but not other types of cerebellar neurons

While the prior data suggest that the neuronal ectopias observed in *Chn2^−/−^* mutants contain *Chn2* expressing cells, we sought to more thoroughly examine the composition of these ectopias. To test for the presence of mature GCs, we made use of the marker Gamma-Amino Butyric Acid Receptor subunit α6 (GABARα6) (Fig. 3a,b). Most cells in the neuronal ectopias in *Chn2^−/−^* animals colabeled with GABARα6, confirming the presence of mature, fully differentiated GCs (71 ± 5%, mean ± SD). Furthermore, we did not detect the immature GC marker Pax6 in the ectopias (Supplementary Fig. S1a,b). To explore the possibility of other cell types contributing to the composition of these ectopic clusters, we immunolabeled with antibodies raised against the Purkinje Cell marker Calbindin, but did not find any Calbindin+ cells within the clusters (Fig. 3c,d). Interestingly, Purkinje cell dendrites failed to invade the space occupied by the neuronal clusters (Fig. 3d). We also immunolabeled for the general GABAergic interneuron marker Parvalbumin (Fig. 3e,f) and found no co-labeling in the neuronal ectopias. Finally, we immunolabeled with the GABAergic marker Glutamic Acid Decarboxylase 67 (GAD67) (Fig. 3g,h). No GAD67^+^ cell bodies were detected in the ectopic clusters. We did detect evenly spread ML labeling of GAD67-positive processes, even in areas containing neuronal ectopias, suggesting these ectopias could potentially receive GABAergic input from stellate or basket cells (Fig. 3g,h). Collectively, these data suggest that the neuronal ectopias found in *Chn2^−/−^* animals are composed primarily of GCs, but not other cerebellar neuronal types.

**Figure 3:**
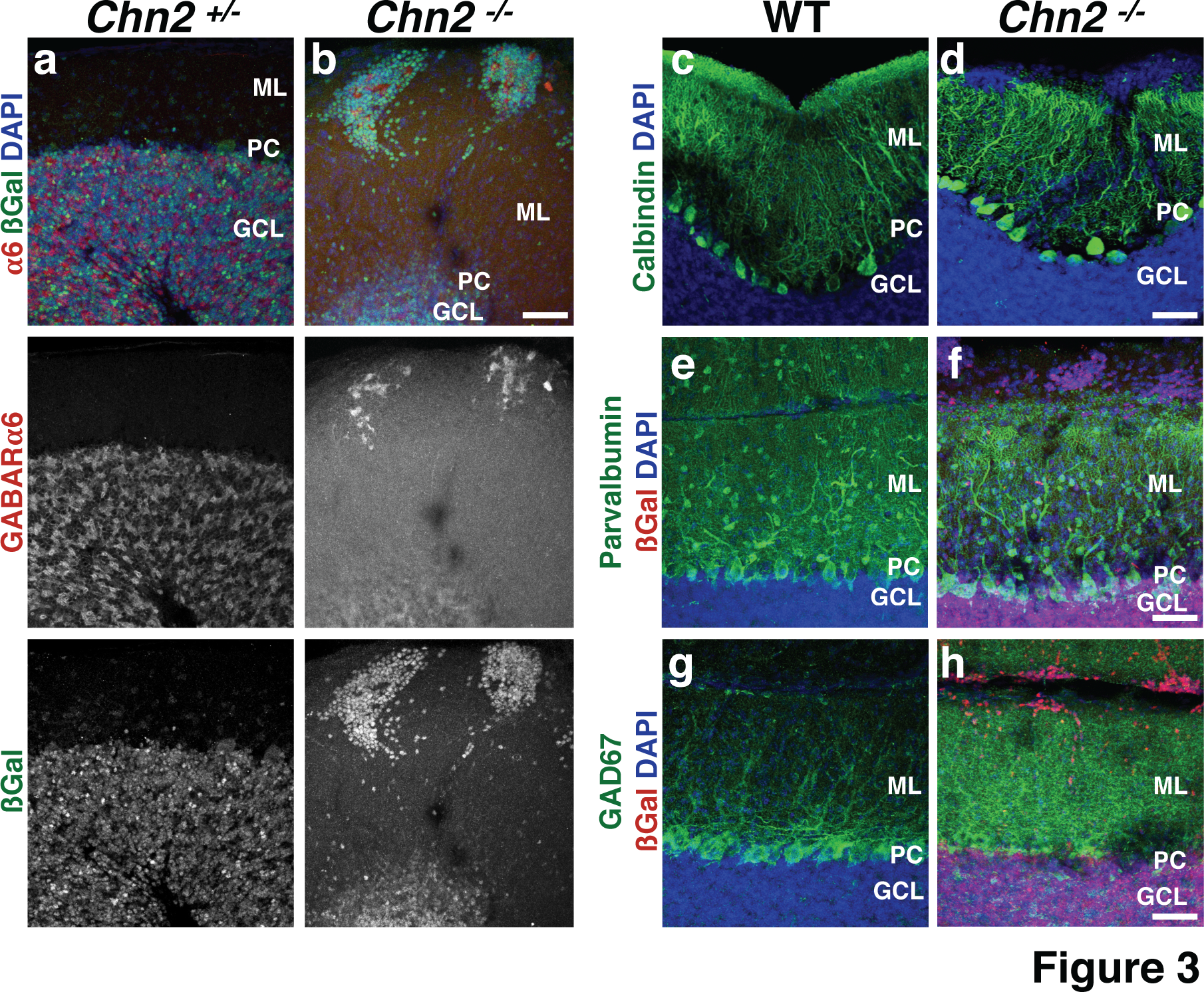
Ectopic clusters contain mature granule cells, but not other types of cerebellar neurons. **(a, b)**Immunofluoresence of the mature granule cell-specific marker GABARa6, with βgal and DAPI as counterstains. βgal positive ectopias contain large numbers of differentiated granule cells. (**c-h**) Immunofluorescence for the PC marker Calbindin (c, d) and the general interneuron markers Parvalbumin (e,f) and GAD67 (g,h), with βgal and DAPI as counterstains. Neuronal ectopias do not co-label with any of these three markers and therefore do not contain PCs, stellate, or basket cells that normally occupy the ML. Interestingly, Purkinje cell dendrites appear to avoid invading the clusters. Scale bar, 50μm for all.

To further study the granule cell migration defect in *Chn2^−/−^* animals, we performed pulse labeling of migrating cells with Bromo-deoxy-Uridine (BrdU), a thymidine analog that incorporates specifically into cells in the S-phase of mitosis. WT and *Chn2^−/−^* animals were injected with BrdU at P10 and cerebella were collected at adult stages. Whereas BrdU labeling in WT animals under these conditions is restricted to the GCL and a few cells scattered in the ML, *Chn2^−/−^* animals display considerable accumulation of BrdU+ cells in the ectopias (Fig. 4a, b). This arrest of GCs born after P10 in the ML of *Chn2^−/−^* mutants suggest that GC migration within the EGL or from the EGL requires *Chn2.*

**Figure 4:**
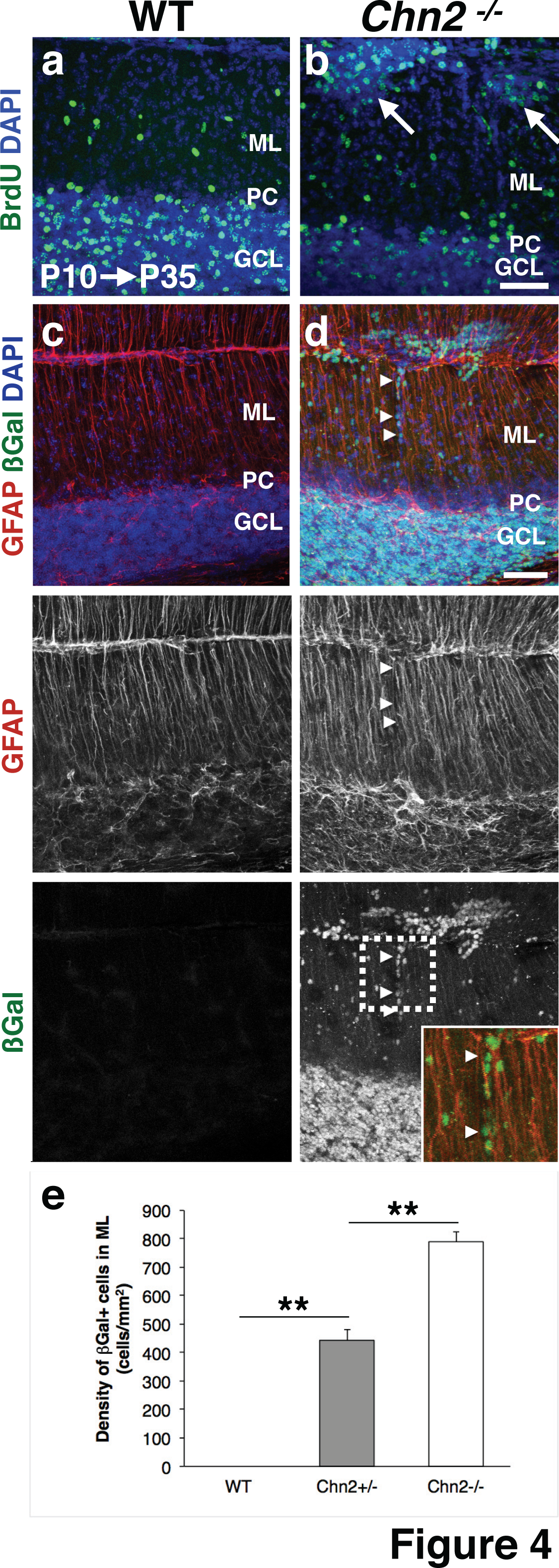
Arrested migration of GCs in *Chn2* null animals. **(a, b)**Cerebellar BrdU pulse labeling at P10, collected at P35. BrdU+ cells accumulate in the ectopic clusters present in *Chn2^-/-^* mutants (white arrows). (**c, d**) Immunofluorescence for βgal and the glial cell marker GFAP, with DAPI as counterstain. The Bergmann Glial scaffold, which radially migrating GCPs adhere to during their migration from the iEGl to GCL, does not appear disrupted in *Chn2^−/−^*mutants. Of note, βgal immunoreactive cells may be seen collected on individual glial tracts (white arrowheads), suggesting some β-chimaerin deficient GCs may initiate but fail to complete radial migration. Insert shows a higher magnification view of the dotted area (green: βgal; red: GFAP). Scale bars, 50μm. (**e**) Quantification of βgal^+^ cells arrested in the molecular layer of WT, *Chn2^+/−^* and *Chn2^−/−^* adult mice. One-way ANOVA, p=1.1102e-16, n=10. **p<0.01, Tukey HSD test. Error bars represent SEM.

During radial migration, GCPs in the iEGL migrate along Bergmann glial fibers to navigate toward the GCL ^4^ Failure of GCPs to properly associate with glial tracts, or errors in glial scaffold architecture itself could inhibit GC radial migration, and could explain the ectopic phenotype observed in *Chn2^−/−^* mutants. Therefore, we examined the structure of the glial scaffolds surrounding ectopic clusters using an antibody raised against Glial Fibrillary Acidic Protein (GFAP) (Fig. 4c, d). We observed no gross alterations to Bergmann Glial structure, arguing against the possibility of an architectural cause underlying the phenotype. However, upon co-labeling with βgal, which strongly marks most cells in neuronal clusters (Fig. 2e), we can observe many individual cells clinging to single GFAP-positive tracts even in the adult (Fig. 4d, white arrowheads). We observe a significant increase in βgal+ cells arrested in the molecular layer of *Chn2^+/−^* and *Chn2^−/−^* animals (Figure 4e; Tukey HSD p<0.01 for WT vs. *Chn2^+/−^,* WT vs. *Chn2 ^−/−^* and *Chn2^+/−^* vs*Chn2^−/−^).* This observation reinforces the idea that GCPs lacking β-chimaerin function stall during radial migration.

### Granule cell ectopias recruit presynaptic partners

In the mature cerebellar circuit, granule cells in the GCL receive glutamaergic input from mossy fibers originating from the spinal cord, pontine nucleus, and other CNS regions. GCs in turn provide glutamatergic input via parallel fibers onto local purkinje cell dendrites ^1^. Since the neuronal ectopias contain differentiated, GABARα6-positive GCs (Fig. 3b), we asked if they could form local circuits. We assayed for the expression of the synaptic marker vesicular glutamate transporter 2 (Vglut2), which labels a subset of cerebellar glutamatergic synapses formed by climbing fibers and mossy fibers, and found robust colabeling with βgal-positive cells within neuronal ectopias (Fig. 5a,b). Furthermore, Vglut2 staining in the ectopias displayed a pattern highly reminiscent of the rosette structures formed by mossy fiber terminals with full penetrance and expressivity (100% of the ectopias displayed this pattern).

**Figure 5:**
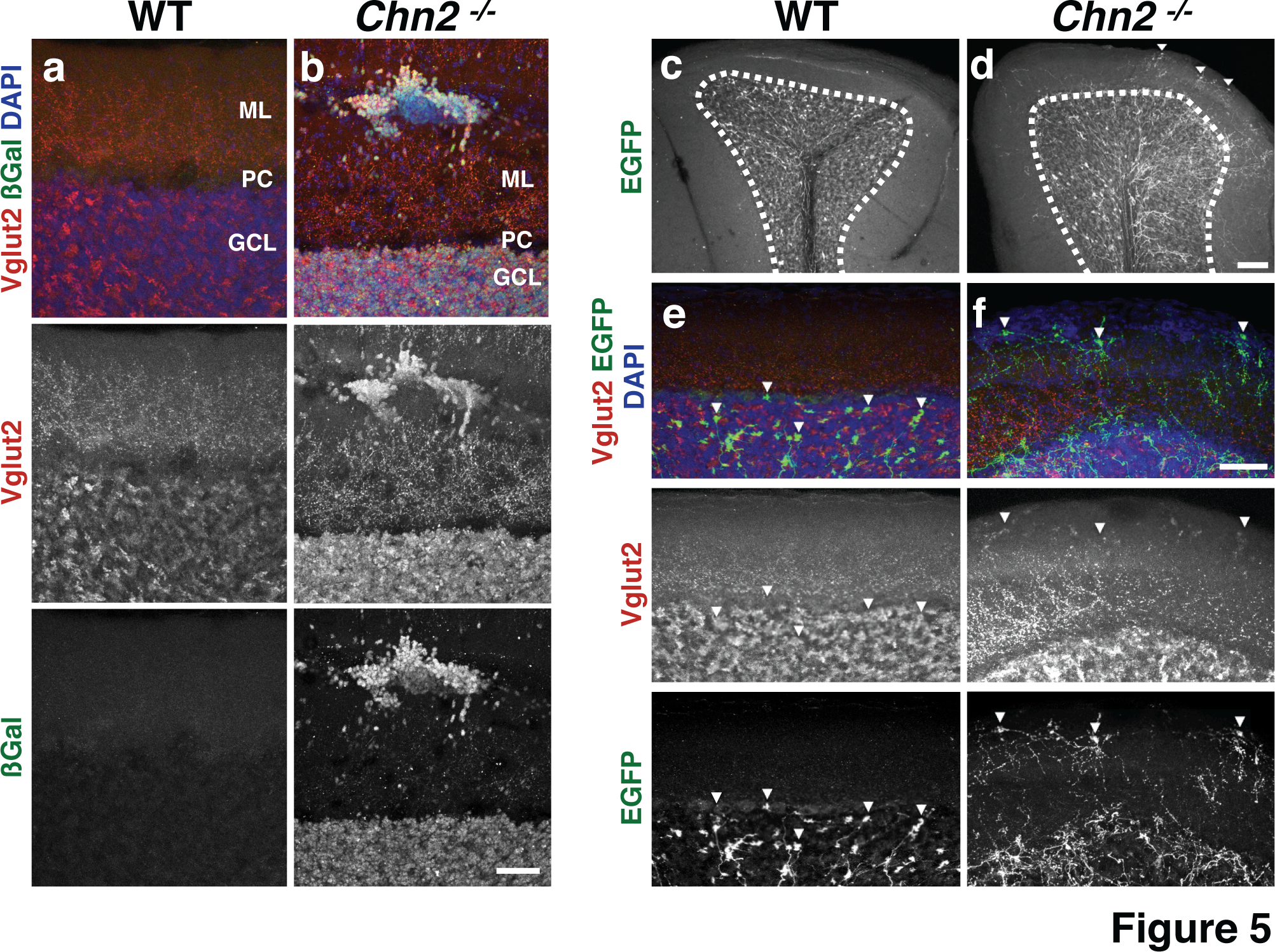
Neuronal Ectopias are contacted by pontine mossy fibers. **(a, b)**Immunofluorescence of the presynaptic marker glutamate vesicular transporter (Vglut2), and βgal, with DAPI as counterstains. 100% of the ectopias robustly label with Vglut2, suggesting they may form synapses with mossy fibers. Scale bar, 50μm. (**c, d**) Injection of AAV-Syn-EGFP into the pons reveals that neuronal ectopias are innervated by aberrant mossy fibers. AAV-syn-EGFP injections into the pontine nucleus of adult (P35) animals label mossy fibers innervating the cerebellar cortex. We found that mossy fibers improperly projected into the ML in β-chimaerin deficient animals (white arrowheads). Scale bar, 100μm. (**e, f**) Higher-resolution image showing that these mossy fibers make direct contact with neuronal ectopias and are surrounded by Vglut2-positive processes. Scale bar, 50μm.

To test whether the Vglut2-positive staining on the ectopic neuronal clusters indeed represented mossy fiber synaptic terminals, we performed stereotactic injections of an Adenosine Associated Virus expressing a Synapsin-promoter-driven Enhanced Green Fluorescent Protein cassette (AAV-Syn-EGFP) into the pontine nucleus. In contrast to control animals, where all observed EGFP-positive axon terminals were restricted to the GCL, we observed EGFP-positive axons extending beyond the GCL in *Chn2^−/−^* mutants (Fig. 5c,d; dotted line demarks outer boundary of the GCL). Furthermore, the EFGP+ axons innervated the ectopias, demonstrating that ectopic GCs could successfully recruit pontine axon fibers. Additionally, under high magnification we found that these terminals co-labeled with Vglut2, suggesting that these represent mossy fiber terminals (Fig. 5e,f).

### External Germinal Layer structure and proliferation is normal in early postnatal *Chn2^−/−^* mice

During cerebellar development granule cells undergo a stepwise maturation process. At embryonic stages, mitotically active granule cell precursors expand across the cerebellar anlage from their point of origin at the rhombic lip to generate the EGL proper. Postnatally, these precursors become postmitotic and extend two horizontal processes, moving inward to generate the inner EGL (iEGL) as a distinct population from the more superficial precursors that remain mitotically active in the outer EGL. In the inner EGL these postmitotic precursors will migrate tangentially, eventually arresting and growing a third perpendicular process. They then begin migrating radially inward, past the PC layer, to form the mature GCL. Given the complex migratory path GCPs take in their development, we asked if an earlier, subtler defect in EGL structure may precede the development of neuronal ectopias.

We examined P10 *Chn2^−/−^* and control animals for the overall distribution of GCPs. We first immunolabeled with antibodies against the transcription factor Pax6, which is active in GCPs in the EGL and maturing GCs in the GCL. We noticed no major difference in Pax6 distribution between *Chn2^−/−^* mutants and controls in the lobules that frequently develop ectopias (Fig. 6 a, b). We also examined the expression profile of the cell adhesion molecule L1-NCAM (L1), which labels migrating granule cells in the inner EGL ^8^. We found no major difference in its distribution between *Chn2^−/−^* mutants and controls (Fig. 6 c, d). These results suggest that there is no altered distribution of GCPs preceding the development of neuronal ectopias. As stated earlier, one possible explanation of ectopia formation is alterations to Bergmann glial tracts. We analyzed the structure of the Bergmann glial scaffold using an antibody against GFAP and found no structural differences in the lobules that more frequently develop neuronal ectopias (Fig. 6 e-f). Collectively, these results suggest that there are no major early postnatal lamination or architectural defects that could predispose certain GCs to arrest.

**Figure 6:**
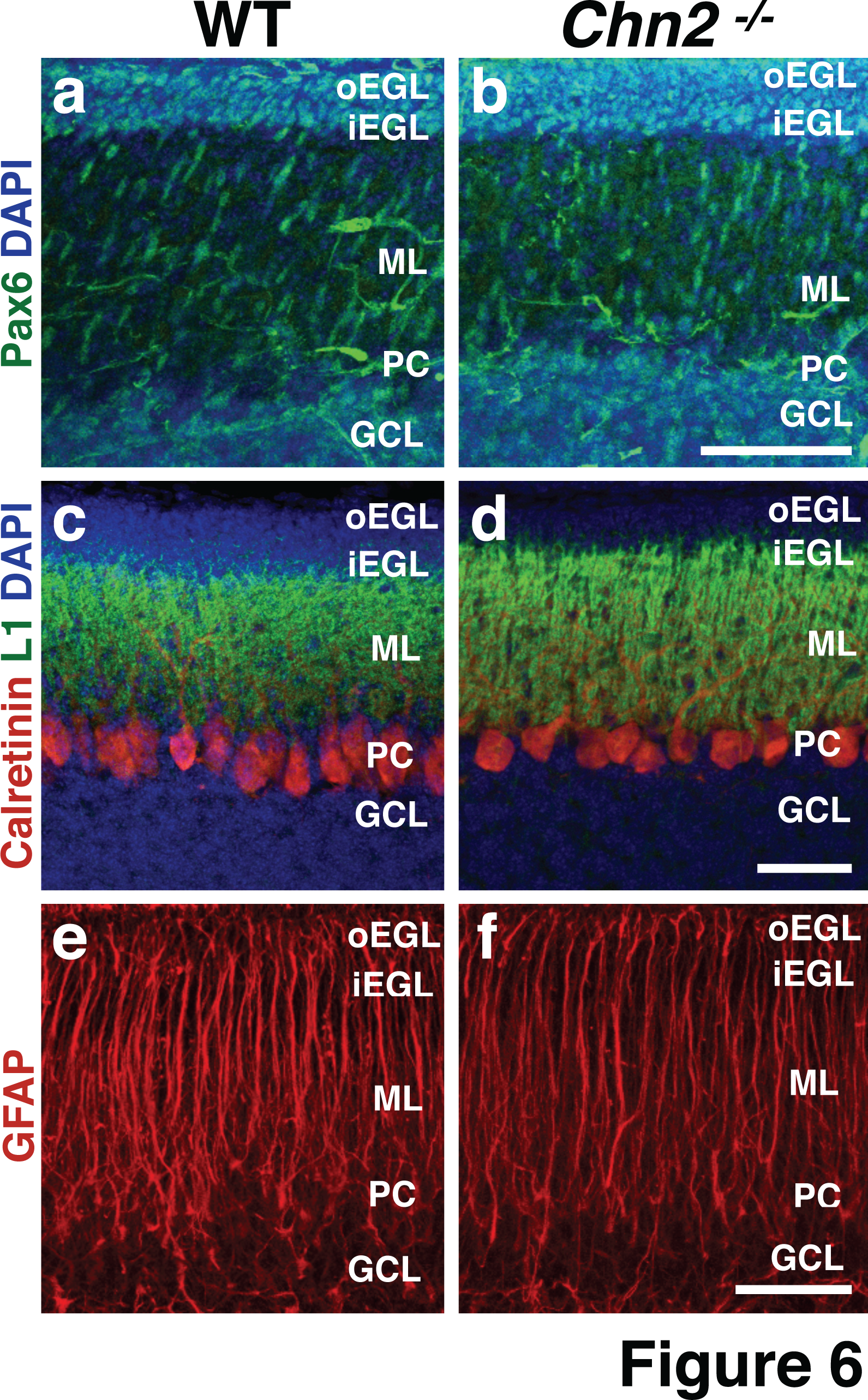
Early postnatal cerebellar structure is unaltered in β-chimaerin deficient animals. **(a-d)**Immunofluorescence on early postnatal (P10) *Chn2^−/−^* mutants with an antibody targeting the transcription factor Pax6, which identifies both GCPs in the EGL as well as GCs in the GCL (a, b) or the cell adhesion molecule NCAM-L1 (L1), which labels migrating GCPs in the iEGL (c, d). At these early postnatal stages, neither Pax6 nor L1 reveal any differences in GCP distribution. (e, f) Immunofluorescence with the glial marker GFAP. The Bergmann glial scaffold appears unaffected. Scale bar, 50μm for a-d; 25μm for e-f.

The data presented in Fig. 4 suggests that in *Chn2^−/−^* mutants GCs are arrested during radial migration. However, defects in tangential migration and/or proliferation could be indirectly contributing to this phenotype. Initial tangential migration of GC progenitors from the rhombic lip appears to be normal in *Chn2^−/−^* animals, as the length of the EGL in WT and *Chn2^−/−^* animals is not significantly different at P0 (Fig. 7a-c; two-tail t-test, p=0.082, n=5) ^29^. A second phase of tangential migration occurs after precursors become postmitotic and move inward to generate the iEGL. It would be predicted that, if GCs are arresting during tangential migration or during the tangential to radial migration switch, the iEGL would become thicker in the folia that develop ectopias in *Chn2^−/−^* mice compared to control animals. To label tangentially migrating GCs in the inner EGL we performed immunostaining with anti-Sema6a antibody at P10 (Fig. 7d,e) ^8,9^ We then measured the thickness of the iEGL relative to the EGL overall in caudal folia (Figure 7f). The iEGL/EGL ratio was comparable in WT and *Chn2^−/−^* mice (Figure 7f; two-tail t-test, p=0.528, n=10), suggesting that tangential migration is not notably disrupted in *Chn2^−/−^* mice.

**Figure 7.**
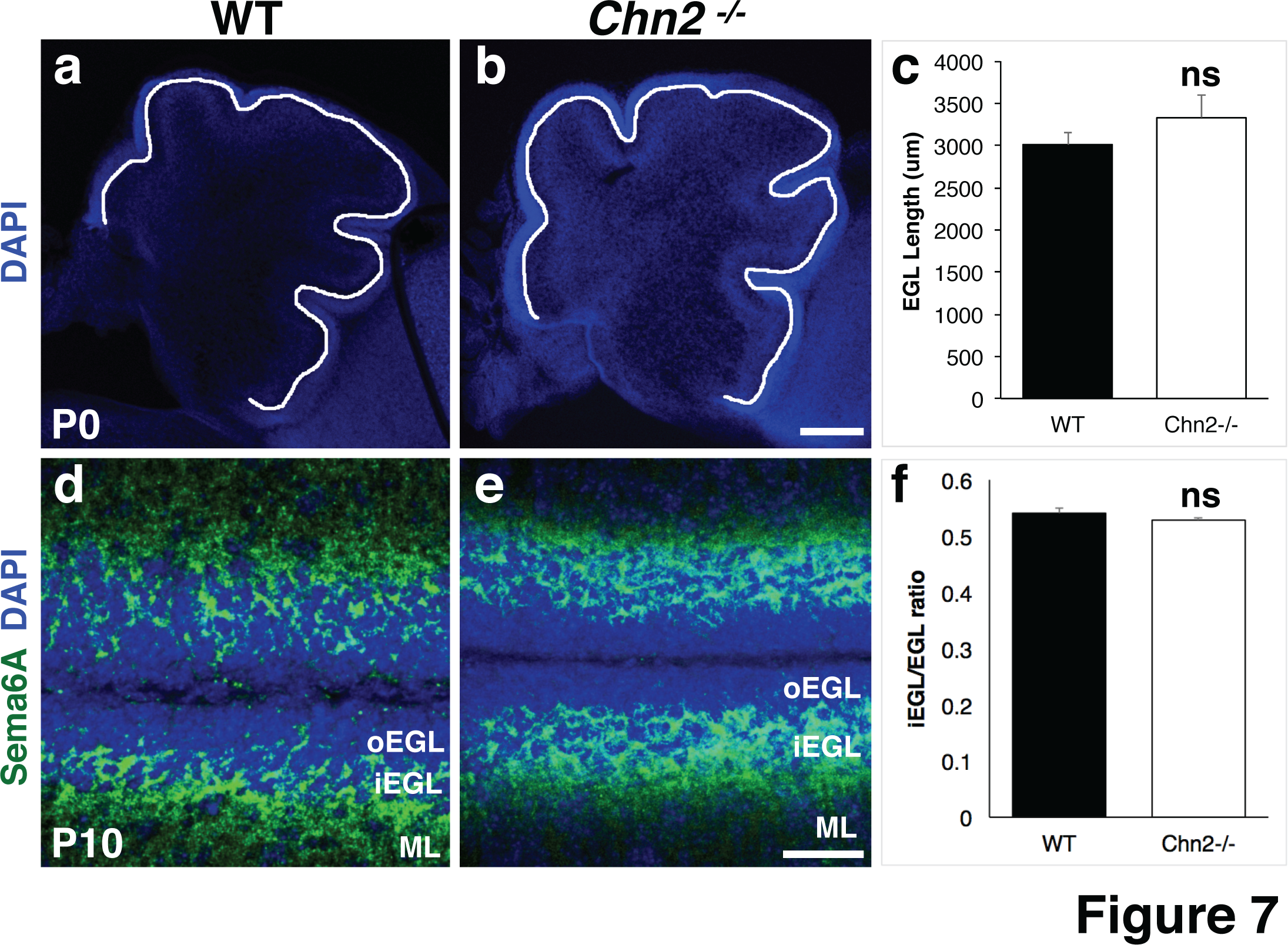
Tangential migration does not appear to be affected in *Chn2^−/−^* mice. **(a-c)**Measurement of EGL length in WT (a) and *Chn2^−/−^* (b) P0 animals. Scale bar, 200μm. (c) Quantification of EGL length. No difference was observed between groups (two-tail t-test, n=5; p=0.082). (**d, e**) Immunostaining for the iEGL marker Sema6A in WT (d) and *Chn2^−/−^* (e) P10 cerebella. Scale bar, 25μm (**f**) Quantification of iEGL (Sema6A+) thickness, relative to overall EGL thickness. Two-tail t-test, n=10; p=0.528. Error bars represent SEM.

During mammalian cerebellar development, granule cell precursors normally continue to proliferate postnatally in the oEGL ^30^. Is proliferation of GCs disrupted during development? We analyzed the distribution of proliferating GPCs in early postnatal animals (P4 and P10) two hours post BrdU injection. Proliferating cells were mainly found in the oEGL in both *Chn2^−/−^* and WT animals (Fig. 8a-f). The density of proliferating GCs in *Chn2^−/−^* and WT animals was comparable (Figure 8c, f; two tail t-test, n=5; P4: p=0.32; P10: p=0.952) suggesting that early postnatal GCPs proliferate normally. To test whether the cells that form the ectopias continue to be mitotically active in the adult, we performed BrdU injections in P35 *Chn2^−/−^* and WT animals (Fig. 8g,h).No proliferating cells were found in the neuronal ectopias, suggesting these ectopic neuronal clusters consist entirely of post-mitotic cells (Fig. 8g,h). Overall these data suggest that the formation of ectopias and the arrest of GCs in the molecular layer of *Chn2^−/−^* animals are primarily due to a defect in radial migration.

**Figure 8:**
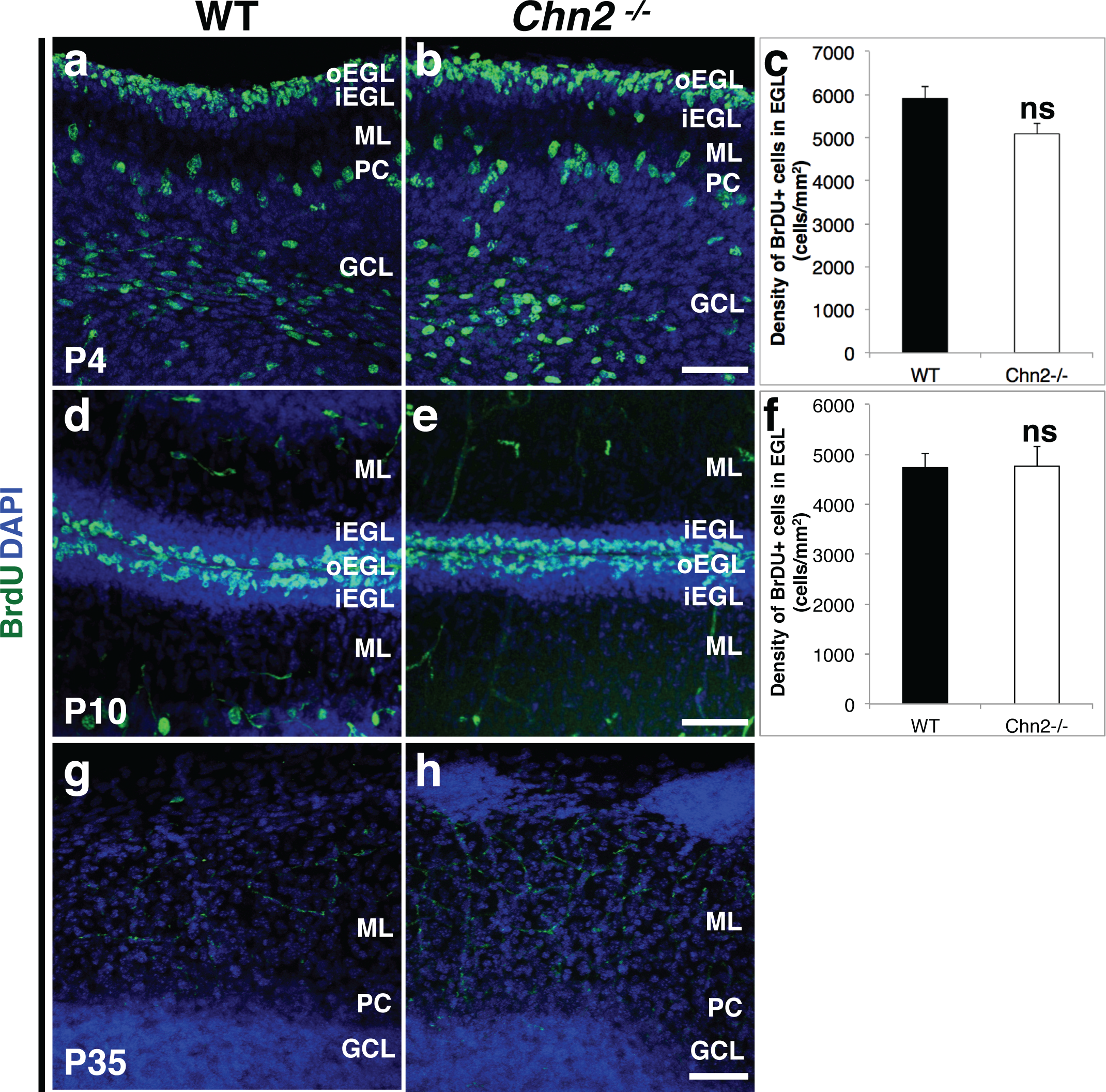
Cell proliferation in *Chn2* deficient animals. **(a-h)**BrdU was injected into either early postnatal stages (P4 and P10, a, b, d, e) or adult (P35, g, h) and allowed to incorporate for two hours prior to animal collection. (c, f) Quantification of BrdU pulse-chase experiments. We found there is no significant difference in the density of proliferating, BrdU-positive GCPs in the oEGL between *Chn2^−/−^* mice and WT animals (n=5 animals per genotype, per stage, 4-5 sections 4μm-thick per animal, two-tail t-test; P4: p=0.32; P10: p=0.952). In adult (P35) animals,neuronal ectopias do not contain proliferating cells, suggesting that they are composed entirely of postmitotic cells (h). Scale bar, 50μm for all. Error bars represent SEM.

### Cerebellar Structure in mice expressing hyperactive β-chimaerin

Genetic ablation of *Rac1* and *Rac3*results in severe disruption of cerebellar granule cell migration ^29,31^. Could increasing β-chimaerin RacGAP activity cause similar phenotypes? To test whether enhanced β-chimaerin activity could also affect cerebellar development, we made use of a knock-in mouse that harbors a hyperactive *Chn2* allele. This allele consists of a single amino acid substitution introduced into the endogenous gene locus ^26^. The I130A substitution yields a protein with a more “open” conformation, which renders it more sensitive to induction ^32^. We collected adult (P35) mice that were homozygous for the hyperactive allele (*Chn2^I130A/I130A^*) and stained for the mature granule cell marker GABARα6 (Fig. 9a, b) and glutamatergic synapse marker Vglut2 (Fig. 9c,d) to label fully differentiated GCs and glutamatergic synapses, respectively. In contrast to *Chn2^−/−^* mutants, *Chn2^I130A/I130A^* animals did not develop ectopic clusters of cells. Further, GC lamination appeared no different from controls. We next looked for other errors in cerebellar structure or lamination by immunostaining for the markers GFAP (Fig.9 e, f), Parvalbumin (Fig. 9g, h), and GAD67 (Fig. 9i, j). We found no difference in the Bergmann Glial scaffold or GABAergic cell populations, respectively. Collectively, these data suggest that hyperactivity of β-chimaerin does not negatively affect cerebellar morphogenesis.

**Figure 9:**
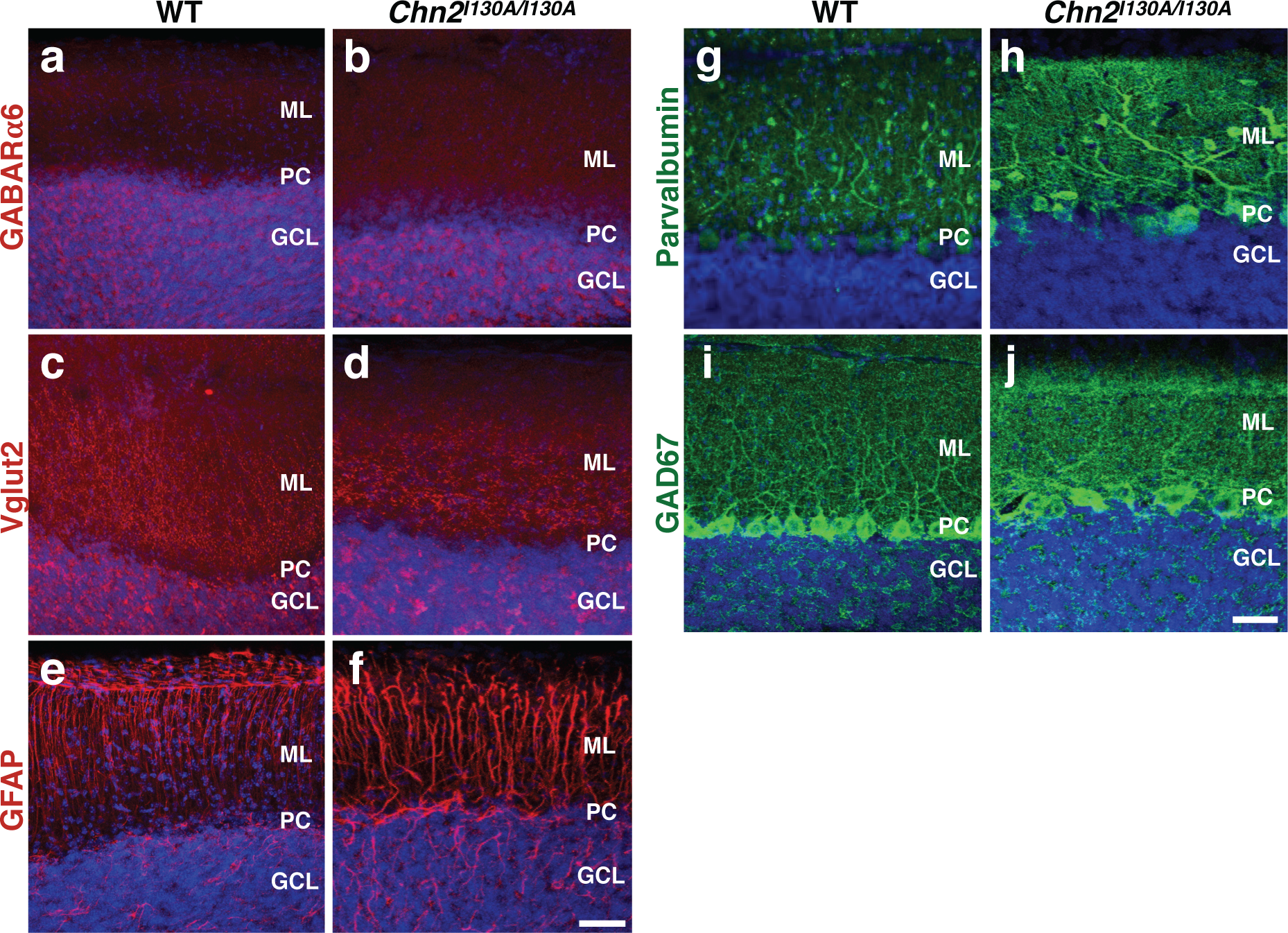
Cerebellar structure is unaffected in β-chimaerin hyperactive mutants. **(a-j)**Histological analysis of adult (P35) cerebellar structure in WT mice and mice homozygous for a hyperactive allele of the β-chimaerin gene *(Chn2^I130A/I130A^*). We observe no notable differences in the mature granule cell marker GABARα6 (a, b), the glutamatergic synaptic marker Vglut2 (c, d), the glial marker GFAP (e, f), or the interneuron markers Parvalbumin (g, h) and GAD67 (i, j). Scale bar, 50μm for all.

## Discussion

Here we show that the RacGAP βChimaerin is essential for cerebellar GC development. Many ligand-receptor pairs have been shown to regulate GC proliferation and migration, but less is known about the cytoplasmic effectors that link these extracellular signals with the cytoskeleton ^5-9^. Guided by the previously reported robust expression of *Chn2* in the adult GCL^27^, we examined whether this cytoplasmic protein could be playing a functional role during cerebellar development. We found that the genetic ablation of *Chn2* results in the formation of ectopic clusters of neurons in the outer ML. These ectopias are primarily formed by GCs. Since we initially established that *Chn2* was mainly expressed in the GCL of early postnatal and adult cerebella (Fig. 1), which represents the mature post-migratory GC population, how could the mispositioned ectopic GCs appear on the outside edge of the cerebellum? Interestingly, a small subset of late pre-migratory GCs expressed *Chn2* mRNA in the outer EGL. Based on the distribution and location of the *Chn2* expressing cells, and the co-localization of βGal with NeuN and α6 in the *Chn2^−/−^* ectopic neuronal clusters, it is likely that the *Chn2^+^* late pre-migratory cells are the ones that fail to migrate inwardly in *Chn2^−/−^* animals. The regulatory mechanisms that restrict *Chn2* expression to a small subset of premigratory GCs are currently unknown, making this a very intriguing question for future studies. The small subset of cells (25-30%) that are part of the ectopias but fail to express the immature GC marker Pax6, the mature GC marker GABARα6, or markers for other neuronal cell types, might represent an intermediate step in GC maturation, as they do express Chn2-driven βGal.

RhoGTPases have been shown to regulate neuronal migration in a variety of neuronal systems ^12,13,15^. In particular, the small G-proteins Rac1 and Rac3 are required for proper granule cell migration ^29,31^. RhoA is also necessary for cerebellar development. To our knowledge, βChimaerin is one of the first RacGAPs to be shown to participate in granule cell migration ^12,15^. Notably, only a small subset of cells in the more caudal cerebellum is affected in *Chn2^−/−^* mutants. Given the essential role of Rac and Rho during cerebellar morphogenesis, other RhoGAPs and GEFs are likely to be involved in regulating these migratory events in other areas of the cerebellum. *Chn1* expression in the developing and adult cerebellum appears to be restricted to the Purkinje Cell Layer ^25,34-36^, making it unlikely for *Chn1* to be performing similar functions as *Chn2* in other cerebellar GCs. With over 80 GEFs and 70 GAPs reported in mammals ^37^, there are many potential additional candidates to be regulating GC radial migration elsewhere in the cerebellum. For example, the Rac GAPs Abr and Bcr have also been shown to participate in GC migration, although they likely act by regulating glial-scaffold development ^38^. While genetic ablation of *Rac1* and *Rac3* reduces the overall level of active Rac, removal of βChimaerin, a RacGAP known to negatively regulate Rac1-GTP levels in neurons ^26,39^, is probably moving the scale in the opposite direction. Thus, balanced Rac activity might be essential for proper GC migration. In this regard, expression of a hyperactive version of βChimaerin (I130A) from the endogenous *Chn2* locus was not enough to disrupt GC migration (Fig. 9). This could be in part due to the regionally and temporally restricted expression of *Chn2* in premigratory GCs.

This novel role of *Chn2* during cerebellar development is the newest addition to a growing list of functional requirements for these RacGAPs during neural development: chimaerins have been shown to regulate axon guidance, pruning in the hippocampus, and cortical lamination ^18-25^. While in the cortex *Chn1* is required for radial migration of most excitatory neurons ^22^, in the cerebellum, *Chn2* is required for migration and positioning of a small subpopulation of GCs, displaying remarkable specificity. The functional requirement of chimaerins during a variety of developmental processes in a wide array of CNS circuits highlights the importance of this small family of RacGAPs during neural circuit formation. Previous studies have demonstrated that Chimaerin function can be modulated by Class 3 semaphorins (Semas) during axon guidance and pruning ^23,26^. In particular, β-chimaerin’s Rac activity was shown to be regulated by Sema3F/Neuropilin 2 function during hippocampal pruning ^26^. Even though cerebellar circuitry proceeds normally in *Sema3F^−/−^* animals^40^, other Semaphorins and Sema receptors have well-established roles during cerebellar GC migration and proliferation ^7-9,41,42^. It is plausible that some of these other members of the Sema family could modulate β-chimaerin function in GCs, since it is still mostly unclear how plexins regulate the actin cytoskeleton ^8,9^

As mentioned above, only a subset of granule cells are susceptible to an arrest in migration in *Chn2^−/−^* cerebella, while the GC population at large is phenotypically normal. Are these ectopic cells able to recruit the right presynaptic partners in a sea of normally positioned GCs? The surprising answer to this question appears to be yes. Anterograde labeling of the pons using viral approaches revealed that the ectopic clusters found in *Chn2^−/−^* cerebella were innervated by pontine axon fibers, one of the normal presynaptic partners for cerebellar GCs (Fig. 5). These ectopic presynaptic terminals are Vglut2+ and display the rosette morphology characteristic of normal pontine mossy fibers. Whether these synaptic terminals are active and mature remains to be explored. While the data presented here provides developmental insight into cerebellar circuit assembly at the anatomical level, it is unlikely that the small number of ectopias present in *Chn2^−/−^* animals will result in obvert behavioral or physiological changes. Far more severe histological defects are observed in other mutant animals without any measurable behavioral or motor changes ^7,8^. Exploration of subtle changes in behavior and physiology in *Chn2^−/−^* animals will be the subject of further studies.

## Materials and Methods

### Animals and Genotyping

The day of birth in this study is designated as postnatal (P) day 0. The generation of *Chn2^−/−^* and *Chn2^I130A/+^* mice has been described elsewhere ^26^. Genotyping of *Chn2^−/−^* mice was performed by PCR using the following primers: *Chn2KO1:* 5’-CAGCCTGGTCTACAGAGTGAG-3’;*Chn2KO2:* 5’-GCATTCCACCACTGAGCTAGG-3’; Chn2KO3: 5’-GTAGGCTAAGCATTGGCTGGC-3’. Genotyping of the *Chn2^l130A/+^* knock-in mice was performed by PCR using the following primers: Chn2KIF: 5’-CCAAGCCCAGCTTTAGAGTGGGC-3’; Chn2KIR: 5’-GAAGGCCCTCCTTTGCTCTGAG-3’. All animal procedures presented here were performed according to the University of California, Riverside’s Institutional Animal Care and Use Committee (IACUC) guidelines. All procedures were approved by UC Riverside IACUC.

### Immunohistochemistry

Mice were perfused and fixed with 4% paraformaldehyde for 2 hours at 4°C, rinsed and sectioned on a vibratome (150 μm). Immunohistochemistry of floating parasagittal cerebellar sections was carried out essentially as described ^43^. The primary antibodies used were: rabbit anti-calbindin (Swant at 1:2500), anti-parvalbumin (Swant at 1:2000), rabbit anti-calretinin (Swant at 1:2000), chicken anti-βGal (AVES labs at 1:2000), chicken anti-GFP (AVES labs at 1:1000), rabbit anti-GFAP (abcam at 1:1000), guinea pig anti-vGlut2 (Millipore at 1:1000), rabbit anti-α6 (Millipore at 1:1000, discontinued), Mouse anti-GAD67 (Millipore at 1:500), rat anti-L1 (Millipore at 1:500) and mouse anti-pax6 (Developmental Studies Hybridoma Bank at 1:200). Sections were then washed in 1 X PBS and incubated with secondary antibodies and TOPRO-3 (Molecular Probe at 1:600 and 1:2000, respectively). Sections were washed in PBS and mounted using vectorshield hard-set fluorescence mounting medium (Vector laboratories). Confocal fluorescence images were taken using a Leica SPE II microscope. Area and length were measured using ImageJ. For cell counts, the ImageJ cell counter plugin was used ^44^

### *In situ* Hybridization

*In situ* hybridization was performed on floating cerebellar vibratome sections (150 μm thickness) using digoxigenin-labeled cRNA probes, essentially as described for whole-mount RNA in situ hybridization ^45^. Generation of the *Chn2* cRNA probes has been described in ^26^

### Injections of AAV

Synapsin-EGFP AAV8 was obtained from the University of North Carolina viral core. The concentrated viral solution (0.2 μl), was delivered into the pons by stereotactic injection (0.25 μl per min), using the following coordinates: anterior-posterior, –5.1 mm; lateral, ±0.6 mm; and vertical, –4.1 mm. For all injections, Bregma was the reference point.

### BrdU labeling

BrdU labeling agent was purchased from Life Technologies (#000103) and was delivered via intraperitoneal injection at 1ml BrdU solution/100g animal weight, following manufacturer instructions. Brains were perfused and collected 2hrs post injection for proliferation assessment, or as adults for pulse-chase experiments. Perfused brains were fixed for 2 hours and sectioned on a vibratome (150 μm thickness). Sections underwent antigen retrieval: incubated in 1M HCl in 1×PBS for 30 mins at room temperature, washed 3×10 min in 1×PBS, incubated in 10mM sodium citrate for 30min at 80C, and washed 3×10min in 1×PBS. Following antigen retrieval, immunohistochemistry was performed as described above using a mouse monoclonal antibody anti-BrdU (Invitrogen, clone BU-1, MA3-071 at 1:250).

## Availability of data and materials

All data analyzed during this study are included in this article.

## Authors Contributions

JAE designed and performed experiments, and wrote the manuscript. WW, YCEW and BML performed experiments. MMR conceived the project, designed and performed experiments, and wrote the manuscript.

## Acknowledgements

We would like to thank Drs. Nils Brose and Andrea Betz, and Dr. Marcelo Kazanietz for sharing with us the Chn2^−/−^ and Chn2^I130/I130^ mice, respectively. We also thank Dr. Kevin Wright for helpful comments on the manuscript. This study was supported by Initial Complementary Funds from the University of California, Riverside

